# Comparing Community Detection Methods in Brain Functional Connectivity Networks

**DOI:** 10.1101/2020.02.06.935783

**Authors:** Reddy Rani Vangimalla, Jaya Sreevalsan-Nair

## Abstract

Brain functional networks are essential for understanding functional connectome. Computing the temporal dependencies between the regions of brain activities of functional magnetic resonance imaging (fMRI) gives us the functional connectivity between the regions. The pairwise connectivities in matrix form correspond to the functional network (fNet), also referred to as a functional connectivity network (FCN). We start with analyzing a correlation matrix, which is an adjacency matrix of the FCN. In this work, we perform a case study of comparison of different analytical approaches in finding node-communities of the brain network. We use five different methods of community detection, out of which two methods are implemented on the network after filtering out the edges with weight below a predetermined threshold. We additionally compute and observe the following characteristics of the outcomes: (i) *modularity* of the communities, (ii) symmetrical node-partition between the left and right hemispheres of the brain, i.e., *hemispheric symmetry*, and (iii) *hierarchical modular organization*. Our contribution is in identifying an appropriate test-bed for comparison of outcomes of approaches using different semantics, such as network science, information theory, multivariate analysis, and data mining.

## 1 Introduction

Understanding the connectivities between different regions in the brain has been a challenge in the area of brain network analysis. Non-invasive and in-vivo imaging techniques are commonly used for brain studies today, owing to the advances in neuroimaging domain. fMRI is one of the widely used brain imaging modalities. Similarly, other modalities such as electroencephalography (EEG) and magnetoencephalography (MEG) techniques are also used to create functional networks (fNet) to analyze the brain activities. The nodes of these networks correspond to regions of interest (ROIs) in the brain confirming to a specific anatomical atlas, e.g., Automated Anatomical Labeling atlas (AAL) [39], Dosenbach atlas (DOS) [12]. The edges between the nodes are computed based on the relationships between all these regions of the brain, which encode the connectivity between the nodes^1^. Here, we focus on the pairwise correlation between nodes in networks computed from fMRI at resting state. For example, the sample network datasets with Brainnet Viewer [42] are computed as correlation matrices. Functional connectivity is inferred from the correlation of the blood-oxygenation level dependent (BOLD) signals of fMRI imaging [40,29] between nodes, as defined for the brain network [38].

In the conventional workflow of brain functional connectivity network (FCN) analysis [13,22,41], these connectivity matrices^2^ are subjected to sparsification by retaining only edge weights of these networks, which are greater than a threshold value. These sparsified matrices are either used directly as weighted graphs or binarized to give unweighted graphs. These preprocessed networks are referred to as *edge-filtered networks*.

Community detection is one of the frequently implemented analysis of FCN. Sporns [37] has discussed about modularity being used for functional segregation and integration, for finding communities and hubs. Functional segregation is the process of identification of ROIs that are related with respect to their neuronal process and are represented as a module. These modules in the network are also referred to as communities, where they have dense intra-community links and sparse inter-community links. Sporns has discussed how functional segregation has been done using multiple approaches, two of which include performing the conventional community detection in the network, and identification of “Resting State Networks” (RSNs), respectively. An RSN is a set of regions in the brain, which show coherent fluctuations of the BOLD signal. Bullmore et al. [6] have described how graph-based methods can be used on brain FCN, and explained the clustering tendency and modular community structure of the brain. In this work, we systematically compare different community detection procedures using an appropriate case study, which is a test bed.

As a complex network with small-world behavior, brain FCN exhibits the property of dense edge connections between nodes of the community and sparse connections across the communities [2]. Meunier *et al.* [25] have discussed how the brain networks, like any other complex networks, have multiple topological scales and hence hierarchical node-groupings, along with modularity. Meunier *et al.* [25] have also explained the existence of both overlapping and non-overlapping communities that display hierarchical modularity. In this work, we focus on non-overlapping node-partitions, *i.e.*, each node belongs to only one module/community. Here, we study the modular behavior of nodes and the hierarchical organization of these modules.

In the edge-filtered networks, network science approaches are strongly influenced by the threshold value used for filtering edges. Since the network topology itself changes drastically depending on the choice of the threshold, the choice has to be carefully made. Jeub *et al.* [18] have used a range of threshold values and a consensus method for clustering the nodes in a completely connected network. Lancichinetti *et al.* [21] have explained the reasons to consider different values of thresholds to get different edge-filtered networks, and then use the consensus of the outcomes from these networks to determine the clusters of a complex network. It is also known that applying a single threshold value on network tends to discard weak and/or negative-signed edges, whose relevance has not been considered [13]. At the same time, finding a threshold interval is also a difficult problem [13]. Given the essential role of edge filtering in FCN analysis, we evaluate its role in community detection by comparing the outcomes using the completely connected brain FCN^3^, *i.e.* without applying a threshold, against the edge-filtered variants of the same network, in a suitable test bed.

### Our Contributions

We compare different functional segregation methodologies on the FCN. The edge-filtered networks reveal the topology of the significant subnetwork(s). However, applying a threshold on the network may not preserve the semantics of the entire network, which calls for independently studying the complete network. We compare the results derived from both edge-filtered and complete networks to evaluate an ideal node partitioning of the given FCN. The crucial questions we address here are:

– How do the chosen approaches implemented on the complete network compare to those on it’s edge-filtered variant?
– Do the different functional segregation methods (tend to) converge at *n* node-partitions, *i.e.*, is there a value of *n* for which the node partitionings tend to be identical?
– If such a number *n* exist, then what is it’s biological significance?
– In different functional segregation methodologies, how can we study the hierarchical organization of the node partitions?

#### Frequently used notations

*Functional Connectivity Network (FCN), Louvain Method (LM), Infomap (IM), Exploratory Factor Analysis (EFA), Hierarchical Clustering (h-clust), Hierarchical Consensus Clustering (HC), Automated Anatomical Labeling atlas (AAL), ground truth (GT).*

## 2 Methods

Our objective is to find the modules in the brain network with maximum modularity, with a preference for methods which extract hierarchical organization within the modules. There are several state-of-the-art approaches for fulfilling this objective. Our gap analysis shows that a systematic comparison of these methods with differences in preprocessing the network is essential to understand the salient aspects of these methods. We select five methods with different underlying principles and where not all use edge filtering, and propose a case study to compare them. Our workflow is given in Figure 1.

**Fig. 1.**
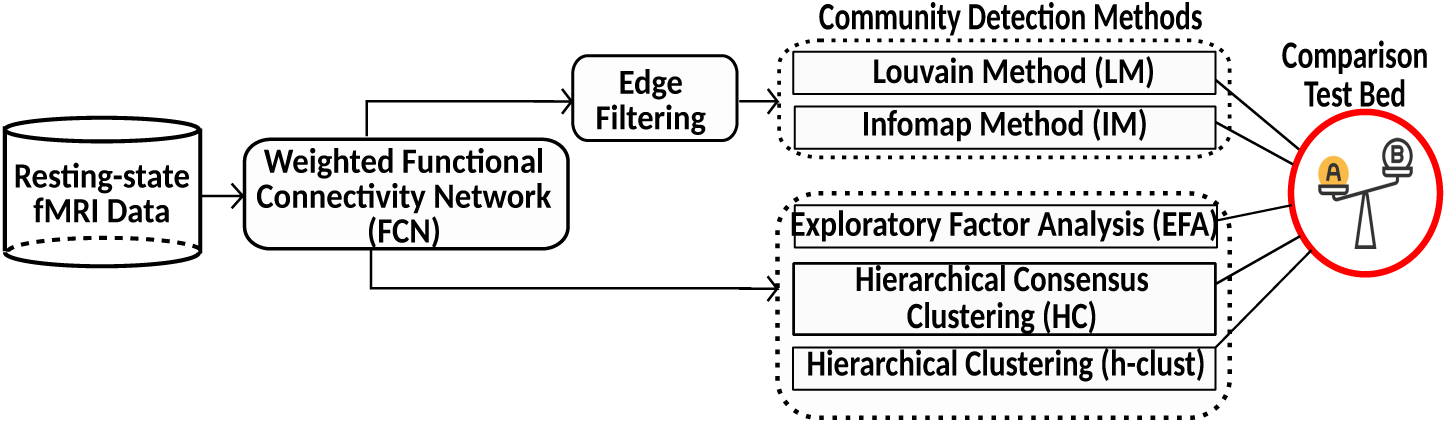
Our proposed workflow for using a test bed for comparing different node partitioning techniques in the human functional connectivity network.

### Network Construction

The FCN is generated using fMRI data from multiple subjects in a cohort. First, an FCN is computed per subject, and the network connects different ROIs, which are the parcellations of the entire brain using a specific atlas, e.g., AAL. The mean time courses (BOLD signal) of the ROIs are extracted, and Pearson’s correlation coefficients are computed between the nodes. Further, Fisher’s r-to-z transformation is applied, thus giving z-score matrices, which are then aggregated across different subjects to get a single un-weighted matrix. Thus, the FCN corresponding to this matrix is a completely connected graph, with the ROIs as nodes and the correlation between them as edge weights. In our work, the choice of the dataset is further restricted by the requirement of positive semi-definiteness of the matrix, so as to make it eligible for exploratory factor analysis (EFA).

### Edge Filtering

Upon filtering out the edges with weights below an appropriate cutoff value [13], the FCN has been shown to exhibit small-world characteristics [23]. Small-world networks have clustering property, which enables finding communities using the modularity measure [14]. Hence, filtering edges is one of the popularly used preprocessing methods in FCN analysis. The threshold for edge filtering is selected by observing a value at which the network changes topology. This change can be identified by analysing statistical properties of the edge weights and their distribution, or by studying the network properties after applying discrete values of threshold, such as node degree distribution (Figure 2(i)) and percolation analysis (Figure 2(ii)).

**Fig. 2.**
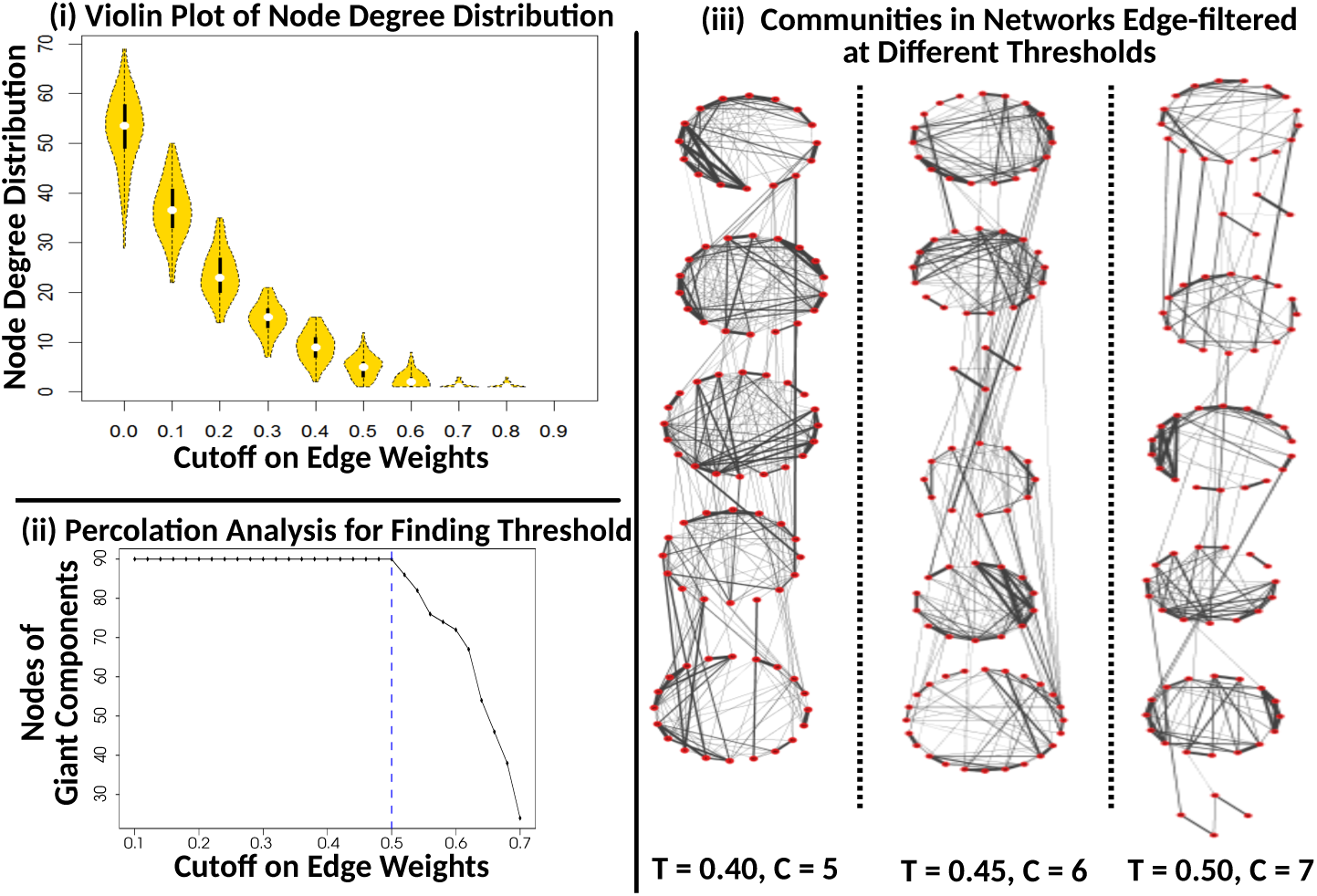
Case study analysis: (i). The violin plot shows the degree distribution of the nodes of the network at different thresholds on edge values, and the elbow curve here is used for finding the optimal threshold for edge-filtering. (ii). The plot of size of giant components (#nodes) at each threshold of edge weights, using percolation analysis [5] is used for finding cutoff. (iii). The networks with edges filtered using different thresholds ‘*T*’ show different topologies, as shown by their graph layout using the stacked circular layout of nodes in their communities ‘*C*’ extracted using Louvain community detection. The edge width is proportional to the correlation value. The plot is generated using Cytoscape, utilizing group attribute layout.

However, applying edge filtering is fraught with stability issues, *i.e.*, slight perturbations in the threshold cause observable changes in the network topology at different threshold values (Figure 2(iii)). The circular layout places the nodes in circles, which correspond to communities. We observe that the network filtered at different layouts show different modular organization. The circular layout, which can be stacked horizontally or vertically, is flexible in showing the instability in network depending on the choice of threshold.

### Node Partitioning

Here, we focus on different node-partitioning methods with non-overlapping communities, which is known as hard clustering. Our objective is to compare five such methods using an appropriate test bed. We use two community detection methods on the edge-filtered network, namely, Louvain community detection (LM) [4] and Infomap (IM) [32]. The remaining three methods, which use the entire correlation matrix, *i.e.*, the complete network, include exploratory factor analysis (EFA) [15], hierarchical clustering (h-clust) [19], and hierarchical consensus clustering (HC) [18]. For the methods used in edge-filtered networks, graph-based techniques automatically provide the number of clusters, which can be used in methods expecting them as inputs.

LM and IM are graph-based methods used for community detection on the sparsified network. LM is a greedy optimization method that maximizes the modularity of the network using an iterative method. Every node is initially considered to be a community, and communities are merged using the nearest neighbor criterion when the modularity value *Q* is computed. The algorithm is iterated until all nodes are grouped with possible maximum modularity value. An information theoretic method, IM is one of the fastest and accurate methods for identifying communities [28] and is widely used in understanding modules in FCN. It is based on the principle that there is a higher likelihood of a random walker most taking steps within a dense community than across communities. The community detection methods LM and IM, essentially exploit the network topology to find appropriate *cuts* in the network to identify densely connected subnetworks. Thus, the methods that are used on edge-filtered network are semantically different from those using the complete network, such as EFA, h-clust, and HC.

EFA is known to be an exploratory or experimental method used for correlation analysis, which uses maximum likelihood function [9] to find *factors*. The factors determine a causal model based on which the correlations between the random variables, *i.e.*, nodes in the FCN here, can be explained. Thus, factors are groups of nodes, which are considered as a node partitioning, modules, or communities, here. h-clust, implemented using different linkage methods, is a clustering technique used in data mining to extract hierarchical clusters. We choose to use h-clust owing to the known structure of hierarchical modularity of the brain FCN [25]. We have experimented with single, complete, average, and ward linkage methods in h-clust. HC method has been exclusively used on brain networks, where the clusters are identified using generalized Louvain community detection [20] method with fixed resolution value (*γ* = 1). The clusterings are aggregated using consensus. Here, we implement HC with 100 clusters and *α* = 0.1 [17], where the parameter *α* decides if co-clustering of two nodes is by chance or by their clustering tendency.

### Modularity

We choose to use the modularity metric, *Q*, to measure the effectiveness of node-partitions of each method. The most widely used Newman-Girvan modularity measure [14,27] is used on both directed and undirected networks, where *Q* measures the difference between the fraction of intra-community edges and the expected fraction of such edges based on node degrees. *Q* is in the range [– 1, 1], where positive values indicate clarity in partitioning. As a first-cut, we do not consider the resolution parameter here.

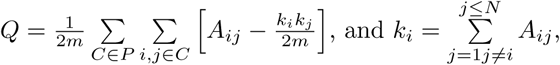

where *A*_*ij*_ is the edge weight between nodes *i* and *j, k*_*i*_ and *k*_*j*_ are degrees of the nodes in the network consisting of *N* vertices, *m* edges, and *C* communities.

### Comparison Test Bed

We propose appropriate settings for comparing the five chosen methods, as there are fundamental differences in the semantics of the methods. We need to ensure that the outcomes are generated with certain fixed settings so that a comparison of the outcomes is scientifically valid. The edge-filtered network used for LM and IM is ensured to be the same. Even though LM and IM automatically give the number of communities, the numbers vary owing to the differences in the methodologies. In EFA, the number of factors *n*_*f*_ is an input. We take the range we have obtained from *n*_*p*_ in LM and IM for *n*_*f*_, so that we can compare the outcomes of EFA with those of LM and IM. For h-clust and HC, since we can use a range of *n*_*p*_ required for different hierarchical levels, we use the same range as is used for EFA.

### Comparative Analysis

We use *Q* for quantitative and Sankey diagrams for qualitative comparisons, respectively. The latter has been used as alluvial diagrams [33] for studying changes in compositions of modules in networks. We additionally perform ground truth analysis, both quantitatively and qualitatively.

## 3 Experiments and Results

We use a specific case study to build the test bed for comparison. We choose a FCN dataset for which we have identified ground truth in literature. We then prepare the edge-filtered variant of the chosen FCN by selecting threshold using different methods. After performing node partitioning using the chosen five methods (Section 2), we perform a comparative analysis of their outcomes.

### Test Bed – Dataset and Ground Truth

We have used the FCN dataset published along with BrainNet Viewer [42], which is generated using the AAL atlas. There are 90 nodes in the FCN. The edge weights are the correlations computed from the resting-state fMRI data of 198 subjects in the Beijing Normal University, provided in the 1000 Functional Connectome Project [3], of healthy right-handed volunteers in the age group of 18-26 years and of which 122 are female. The fMRI scanning was performed in the eyes-closed (EC) state of subjects in state of wakefulness. The network is generated after removing data of one subject owing to rotation error. The test bed requires a ground truth (GT) for this specific dataset, for which we use the findings on a similar dataset used by He *et al.* [16]. Even though the fMRI data in our case study and that identified as GT is different, the demographics of the subjects involved and the processing done on the two datasets are the same. Hence, we take the result of five functional modules by He *et al.* [16] to be the GT, *i.e.* the reference communities. The module identification for the GT has been done using simulated annealing approach, thus, avoiding *similarity bias* with any of our chosen methods.

### Community Detection in FCN

We have compared the communities of the network obtained using five different methods, i.e., LM, IM, EFA, h-clust, and HC, after preparing the test bed (Section 2). We compute an edge-filtered variant of the FCN by identifying an appropriate threshold using the inferences from elbow graph for degree distribution at each threshold (Figure 2(i)) and using percolation analysis (Figure 2(ii)). Since we get optimal thresholds as *T* = 0.4 and *T* = 0.5, respectively, we have checked the *Q* of the communities extracted using LM, and used the one with the higher value of *Q*. At *T* = 0.5, we observe the disintegration of a giant connected component in the network (Figure 2(iii)). When *T >* 0.5, in Figure 2(i), we observe that the node degree distribution is uniform, and the network exhibits uniform topology rather than communities. Overall, at *T* = 0.4, we observe more stability in the dataset; hence, we have chosen *T* = 0.4, as the optimal threshold for the edge-filtered variant to be used in LM and IM. We have also verified against the binarized thresholded version of the network that has been published [42], the threshold used is *T* = 0.4.

We have also run experiments with threshold *T* = 0.45, and *T* = 0.5 to study the change in topology (Figure 2(iii)). We have observed that for *T* = {0.4, 0.45, 0.5}, we get {5, 6, 7} communities using LM, and {7, 9, 12} using IM, respectively. In our case study, IM leads to over-segmentation. We have also observed that for *T >* 0.6 the community detection of the network using LM does not include all the 90 nodes.

The methods on complete networks, namely EFA, h-clust, and HC, require the number of modules as input to give outputs to be compared with those of LM and IM. The optimal value of *n*_*f*_ for EFA is computed using a scree plot [7] and parallel analysis. In our case study, *n*_*f*_ = 9 is the optimal number of factors according to the parallel analysis scree plot. However, we have empirically chosen *n*_*f*_ = 5, given that modularity score is highest for this value, and also, this is equivalent to the GT. Figure 3(i) shows us that for all the methods, the highest *Q* value is observed when the network has five modules, which confirms with the GT. For *n*_*p*_ = 5, LM shows the maximum *Q*, which can be attributed to its greedy characteristic. At five modules, EFA performs at par with LM.

**Fig. 3.**
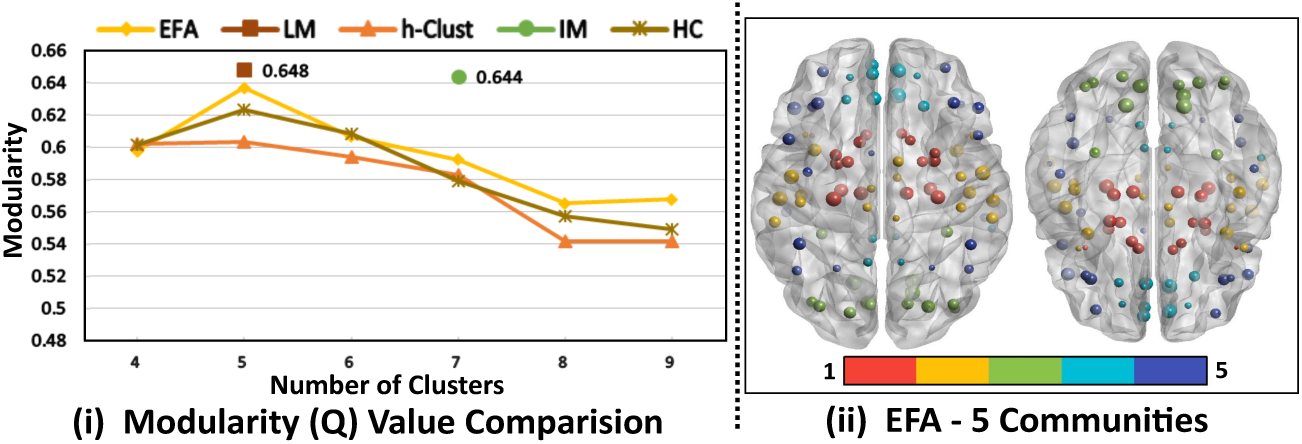
(i). Modularity (*Q*) values of node partitioning for LM, IM, EFA, h-clust with average linkage, and HC with *α* = 0.1, show trends for hierarchical modules, and high values for LM and IM. (ii). Hemispheric symmetry of nodes or ROIs can be observed in the visualization of modules in FCN using brain-surface visualization [42] (BNV), a MATLAB tool.

We use the BrainNet viewer [42] for visualizing the node-communities on the brain surface (Figure 3(ii)) in the spatial context. The axial view of the brain shows *modular organization spatially, i.e.*, neighboring nodes are grouped in a module and *hemispheric symmetry* of the nodes. Hemispheric symmetry implies that both left and right hemispherical nodes of the same brain region tend to co-cluster. EFA with *n*_*f*_ = 5, LM on the network with edge-filtering at threshold *T* = 0.4, and GT demonstrate similar modules, but with modular organization and hemispheric symmetry. We have additionally implemented each of our proposed approaches, independently as ensemble runs, *i.e.* implemented multiple times with slight changes in parameters, *e.g., n*_*f*_ for EFA, and treecut for h-clust and HC. Our motivation is to compare hierarchical modular organization in FCN.

### Comparative Analysis

The Sankey plot [30] or alluvial diagram [33] effectively demonstrates a qualitative comparison of the composition of communities. Figure 4(i) demonstrates that at *np* = *nf* = 5, outputs of LM and EFA are similar, as 83 out of 90 nodes were grouped similarly in both the methods. At *n*_*p*_ = *n*_*f*_ = 5, LM and EFA has the highest modularity value *Q*. The edge crossings in (Figure 4(i)) between LM and EFA are due to one node in the AF4^4^ cluster, and six nodes in the AF5 cluster in EFA. We observe a similar degree of mismatch between EFA with h-clust, at *n*_*f*_ = *n*_*p*_ = 5. However, unlike the mismatch with LM, the community sizes in h-clust are not uniformly distributed as in EFA and LM. In h-clust, we use the consensus of the node-groupings with different linkage methods of hierarchical clustering, namely single, complete, average, and ward. The tree-cut is the deciding parameter for *n*_*p*_, and hierarchy is guaranteed with all linkage methods, by design. The matching is at 86.67%, *i.e.*, 12 out of 90 nodes showed grouping different from EFA. Except for the cluster AH2, the other clusters in h-clust have inconsistent mappings with EFA(Figure 4(i)). When compared to single, complete, average, and ward linkage methods of h-clust, the average-linkage method exhibited the highest matching percentage with EFA.

**Fig. 4.**
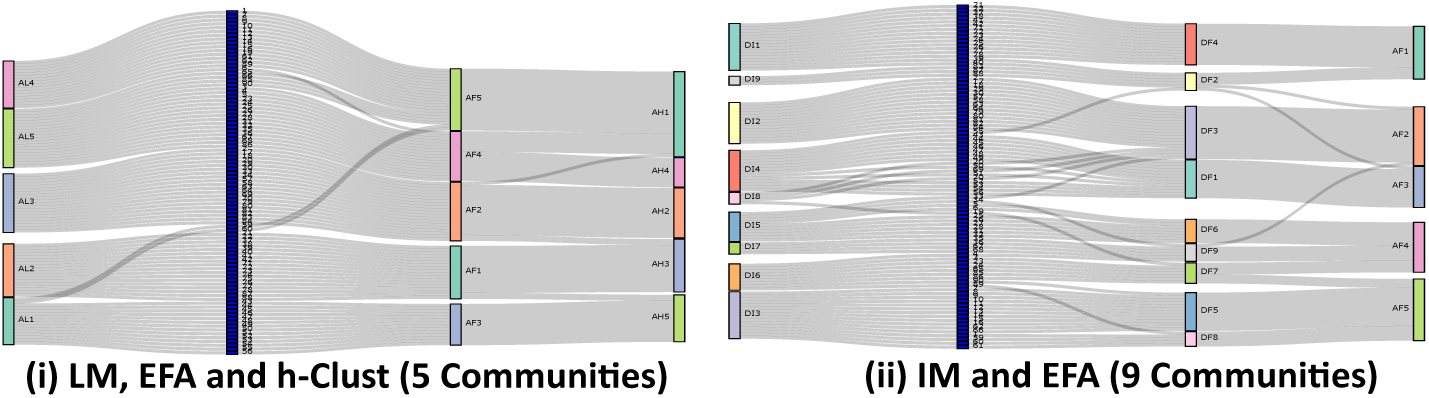
The composition of node-communities from multiple methods is compared using Sankey plot, where the middle vertical bar (in blue) corresponds to the node-IDs, and LM and IM are computed on the edge-filtered network at a threshold *T*. (i). Comparison of LM at *T* = 0.4, EFA at *n*_*f*_ = 5, and h-clust at *n*_*p*_ = 5 (average linkage) LM and EFA shows more similarity of composition and sizes of communities than LM and hclust. (ii) Comparison of IM with *T* = 0.45, EFA with *n*_*f*_ = 9, and EFA with *n*_*f*_ = 5 show differences between IM and EFA, including fragmentation in IM. The naming convention of the communities is given in the footnote^4^.

Interestingly, we have observed that IMon the network with edges filtered at threshold *T* = 0.45 gives nine communities, and the optimal *n*_*f*_ = 9 for EFA, as per the parallel analysis scree plot. Hence, in our test bed, we compare the outcomes between IM (*T* = 0.45) and EFA (*n*_*f*_ = 9). We observe more edge crossings between IM and EFA, indicating that the node-groupings failed to display similar correspondence (Figure 4(ii)). But, interestingly, we observe lesser edge crossings between EFA (*n*_*f*_ = 5) and EFA (*n*_*f*_ = 9) modules. This observation is due to the revelation of the characteristic of hierarchical modularity in the FCN when progressively increasing *n*_*f*_ in EFA.

When comparing against GT [16] using Sankey diagrams as in Figure 5(i) to (iv), we observe that the edge-crossings and inconsistent node-groupings are less in the case of EFA, and more in the case of h-clust. The matching of results with GT is 90.00%, 88.89%, 85.56%, and 83.34%, for EFA, LM, HC, and h-clust, respectively. We observe that the edge crossings are least in the case of EFA-GT-LM, for *n*_*f*_ = *n*_*p*_ = 5 (Figure 5(i)) and HC-GT-h-clust, for *n*_*p*_ = 9 (Figure 5(iv)), indicating similar grouping. But when we closely observe in the latter, the size distribution of the communities in HC-GT-h-clust does not match.

**Fig. 5.**
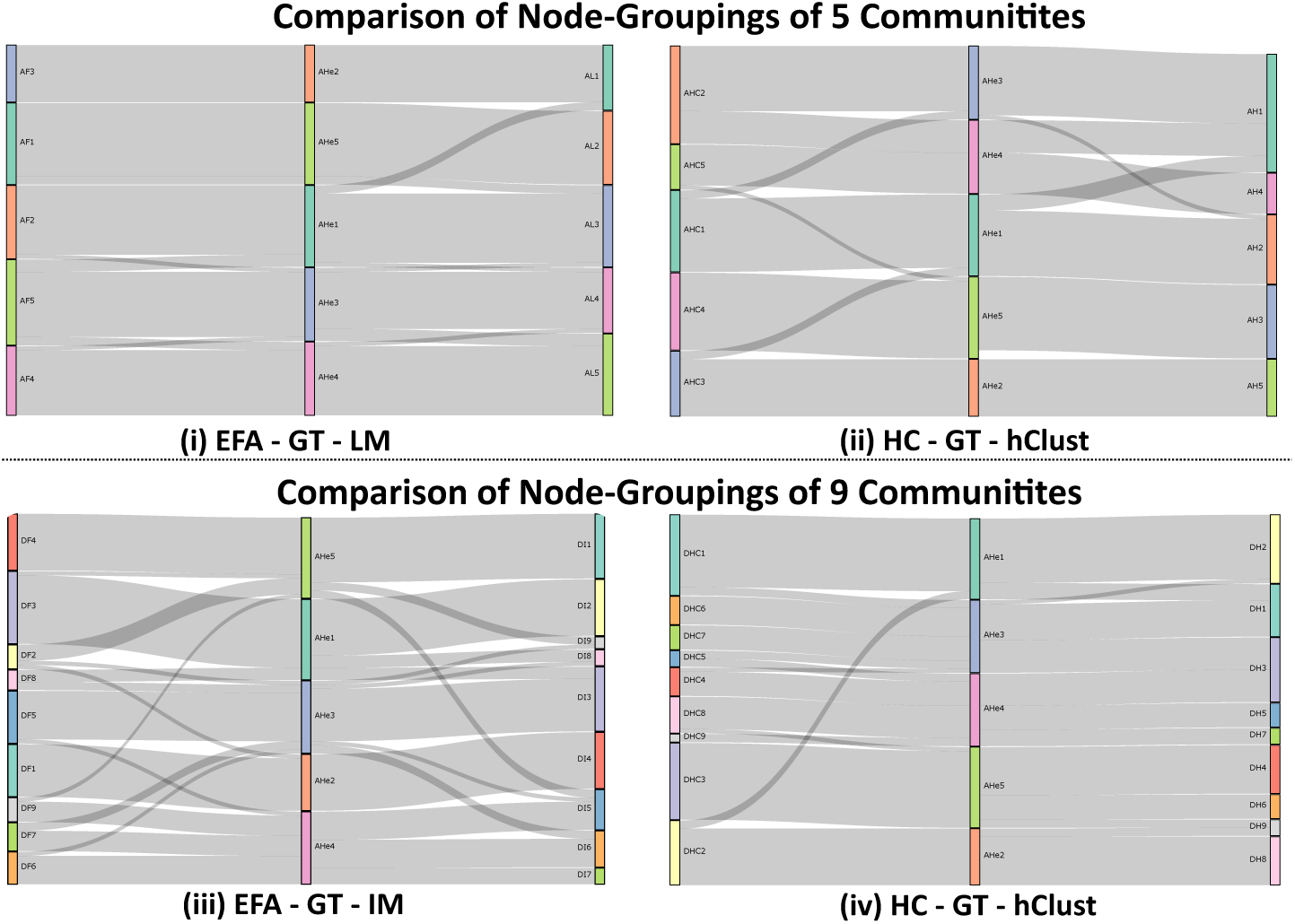
Comparative visualization of mapping of nodes between our selected approaches and ground truth (GT) in [16]. Comparison against GT of (i). EFA (*n*_*f*_ = 5) and LM (*T* = 0.4), (ii). h-clust and HC, at *n*_*p*_ = 5, (iii). EFA (*n*_*f*_ = 9) and IM (*T* = 0.45), (iv). HC and h-Clust at *n*_*p*_ = 9. The naming convention of the communities is given in the footnote^4^.

Overall, we conclude that when *n*_*f*_ = *n*_*p*_ = 5, EFA and LM, with *T* = 0.4, behave similar to each other, and also with GT. Additionally, EFA exhibits hierarchical modularity in our case study.

### Hierarchical Modularity

We know that h-clust and HC show hierarchy in their community detection or clustering owing to the design of preserving hierarchy by performing divisive hierarchical clustering. However, we observe the same in EFA, when we increase or decrease *n*_*f*_ within the range [5, 9]. From Figure 5, we have observed how EFAconfirms with the GT, and now we observe the hierarchical characteristic. Thus, in this case study, we observe that outcomes of EFA demonstrates a hierarchical modular organization of the functional brain. In Figure 6(i), module AF1 is subdivided into BF1 and BF6 when transitioning from 5 factors or 6 factros; similarly, module BF4 has subdivided into modules, CF4 and CF7, to grow from 6 factors to 7 factors, and a similar pattern can be observed when transitioning from 7 to 9 clusters.

**Fig. 6.**
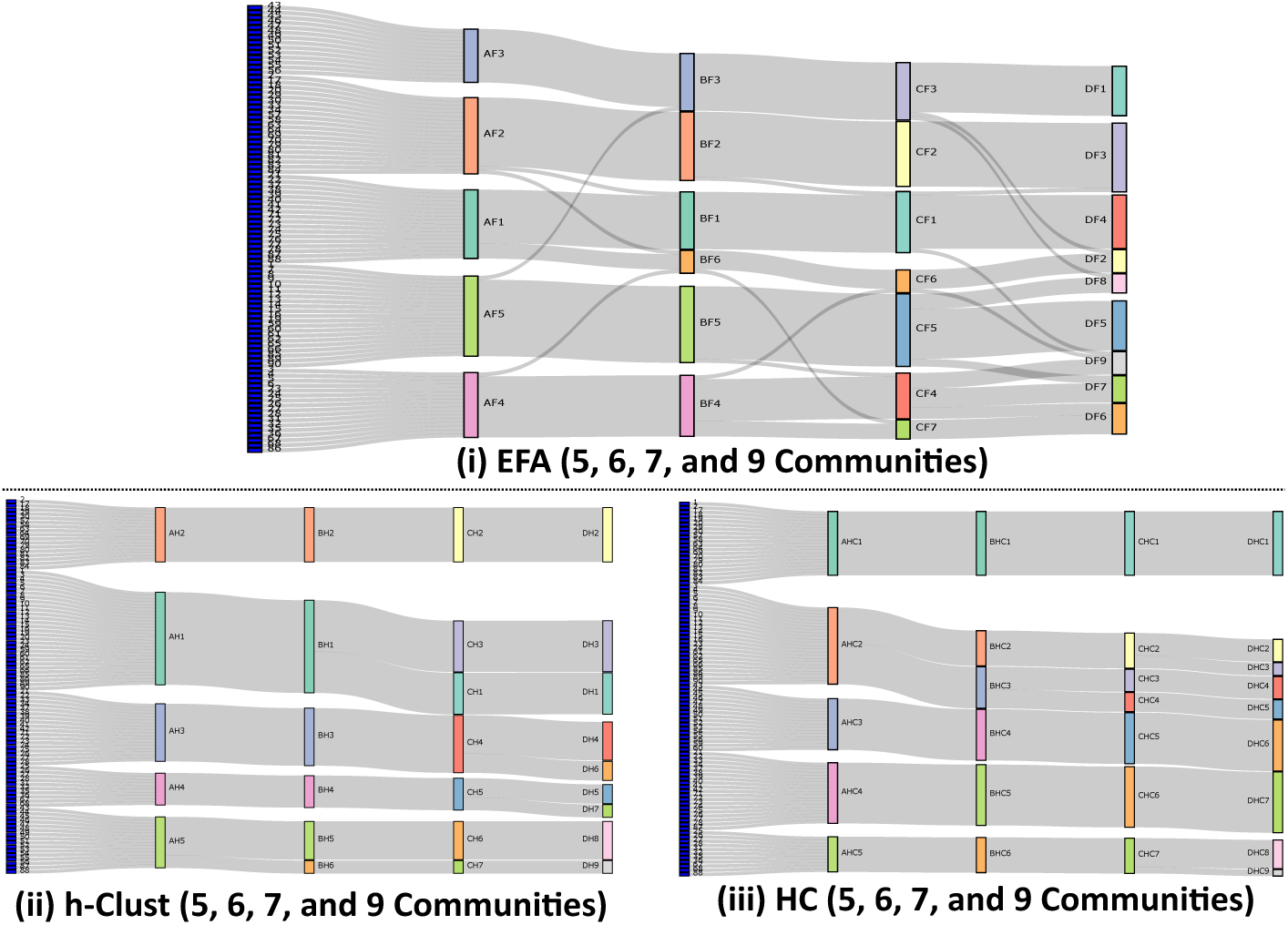
Visualizing the hierarchical modularity from 5 to 9 modules using a cascading effect using Sankey plot using communities identified using (i). EFA, (ii). h-clust, and (iii). HC. The naming convention of the communities is given in the footnote^4^.

We observe the hierarchical modularity of the network when using h-clust, and HC (Figure 6(ii) and (iii)), as per design, when using tree-cut for deciding parameter for module identification. However, both h-clust and HC failed to get outcomes similar to GT, thus violating the hemispheric symmetry of modules.

### Discussions

Our analysis gives results similar to that of Mezer *et al.* [26], where node communities exhibited *symmetric* patterns between left and right hemispheres. We observe hemispheric symmetry in Figure 3(ii), and 4(i), where EFA showed both better symmetry and hierarchical modularity. For example, sensorimotor and auditory regions of the first group in EFA (*n*_*f*_ = 5) are sub-divided into two different groups as the sensorimotor and auditory regions in EFA (*n*_*f*_ = 6). Similar studies of resting-state fMRI [8,43] have confirmed node-communities of the functional connectivity network are symmetrically organized between homotopic regions. Similarly, the modules identified using Newman’s modularity algorithm in [24] has a similar grouping of nodes, with the communities we got from LM and EFA. The regions, hippocampus, and thalamus of sensorimotor system, have always exhibited symmetric patterns between the left and right hemispheres and are consistently clustered together in the same group for all values of *n*_*p*_ and *n*_*f*_.

The biological significance of these five modules is that they correspond to largest functional modules known to exist in the brain [25] as well as the resting state networks usually extracted from fMRI data [36]. The five modules are medial occipital, lateral occipital, central, parieto-frontal and fronto-temporal systems.

Such comparisons would not have been possible without establishing our proposed test bed. We have demonstrated that an appropriately designed test bed enables comparison of outcomes of module identification across different methodologies governed by different semantics. There are shortcomings in our methodology related to: (i) identification of thresholds for edge filtering, as well as (ii) comparing properties of the methodologies. While identifying thresholds, specific discrete values, *i.e.* {0.4, 0.45, 0.5}, were considered which could have missed values in the intervals. A better idea would be to use ranking of edge weights in the network to eliminate one edge at a time, and then analysing the network. In this work, we have not compared the robustness of our chosen methods and properties related to reproducibility [16].

## 4 Conclusions

In this work, we have compared five different approaches for community detection of functional connectivity network. Firstly, using edge-filtering for topological analysis of functional brain networks, we have chosen node-community detection as an outcome of our proposed workflow. In lines of node-community detection, we have proposed the use of graph-based Louvain and information-theoretic based Infomap methods on edge-filtered weighted networks. Secondly, we have introduced matrix-based exploratory factor analysis, distance-based hierarchical clustering and hierarchical consensus clustering for node-community detection, based on the semantics of the connectivity matrices. Using our proposed test bed, we can now answer the question: “For the chosen approaches and an appropriate set of parameters for implementing them, does a number, *n*, exist for a specific FCN, such that the node partitionings tend to be identical?”. The answer is **five** for the dataset we have worked on.

Our work demonstrates how a test bed can be used for systematic comparison of community detection methods in FCN. We have also shown how an explorative method, such as EFA, which is semantically a correlation analysis method, can be used for community detection. Our observations are that both Louvain community detection and EFA perform equivalently when extracting the optimal number of communities on the network. EFA additionally showcases hierarchical organization, when changing *n*_*f*_ progressively. Our work assesses the biological significance of our findings against the GT. Thus, overall, in our case study, EFA performs well in ground truth analysis, and characteristics of hemispheric symmetry of nodes in FCN and hierarchical organization.

However, in the context of newer trends in the functional connectivity studies, our study on resting-state data can be extended to specific cognitive task-based studies. Our study is valuable as a first step towards the comparative analysis of node-partitioning in FCNs across networks with different preprocessing steps.

## ACKNOWLEDGMENTS

This work has been supported by the Visvesvaraya PhD Scheme for Electronics and IT, the Ministry of Electronics and Information Technology, Government of India. The authors acknowledge the use of third-party packages, namely, R packages, including ‘igraph’, ‘psych’ and ‘GPArotation’ for implementing node grouping, [11,31,1,35,10], Cytoscape [34], and BrainNet Viewer [42] for the implementation of the proposed work.

## Footnotes

1 Connectivity matrices of functional networks could be computed using several methods [13], e.g., correlation, mutual information, phase coherence.

2 Connectivity matrix of a **network** corresponds to the adjacency matrix of the **graph**.

3 We refer to these networks as the *complete network*.

4 In Figures 4, 5, and 6, the communities are named in the format *XY* where *X* is {A, B, C, D}, which corresponds to {5, 6, 7, 9} node-communities, and the value of *Y* is {L, I, F, H, HC, He}, which corresponds to {LM, IM, EFA, h-clust, HC, GT}, respectively. For example, **AL4** represents the fourth community out of five communities (*A = 5*) identified using the method LM (*L*).

